# A Type I Restriction-Modification System Associated with *Enterococcus faecium* Subspecies Separation

**DOI:** 10.1101/408914

**Authors:** Wenwen Huo, Hannah M. Adams, Cristian Trejo, Rohit Badia, Kelli L. Palmer

**Affiliations:** Department of Biological Sciences, University of Texas at Dallas, Richardson, Texas 75080

**Author notes:** Contact information for corresponding author: Kelli Palmer.

**Keywords:** Restriction-Modification, *Enterococcus faecium*, genome defense

## Abstract

The gastrointestinal colonizer *Enterococcus faecium* is a leading cause of hospital-acquired infections. Multidrug-resistant (MDR) *E. faecium* are particularly concerning for infection treatment. Previous comparative genomic studies revealed that subspecies referred to as Clade A and Clade B exist within *E. faecium*. MDR *E. faecium* belong to Clade A, while Clade B consists of drug-susceptible fecal commensal *E. faecium*. Isolates from Clade A are further grouped into two sub-clades, A1 and A2. In general, Clade A1 isolates are hospital epidemic isolates whereas Clade A2 isolates are isolates from animals and sporadic human infections. Such phylogenetic separation indicates that reduced gene exchange occurs between the clades. We hypothesize that endogenous barriers to gene exchange exist between *E. faecium* clades. Restriction-modification (R-M) systems are such barriers in other microbes. We utilized bioinformatics analysis coupled with second generation and third generation deep sequencing platforms to characterize the methylome of two representative *E. faecium* strains, one from Clade A1 and one from Clade B. We identified a Type I R-M system that is Clade A1-specific, is active for DNA methylation, and significantly reduces transformability of Clade A1 *E. faecium*. Based on our results, we conclude that R-M systems act as barriers to horizontal gene exchange in *E. faecium* and propose that R-M systems contribute to *E. faecium* subspecies separation.

**IMPORTANCE:** *Enterococcus faecium* is a leading cause of hospital-acquired infections around the world. Rising antibiotic resistance in certain *E. faecium* lineages leaves fewer treatment options. The overarching aim of the attached work was to determine whether restriction-modification (R-M) systems contribute to the structure of the *E. faecium* species, wherein hospital-epidemic and non-hospital-epidemic isolates have distinct evolutionary histories and highly resolved clade structures. R-M provides bacteria with a type of innate immunity to horizontal gene transfer (HGT). We identified a Type I R-M system that is enriched in the hospital-epidemic clade and determined that it is active for DNA modification activity and significantly impacts HGT. Overall, this work is important because it provides a mechanism for the observed clade structure of *E. faecium* as well as a mechanism for facilitated gene exchange among hospital-epidemic *E. faecium*.

## INTRODUCTION

*Enterococcus faecium* is a Gram-positive opportunistic pathogen that normally resides in the gastrointestinal tracts of humans and other animals (1, 2). *E. faecium* can cause life-threatening infections such as endocarditis and is among the leading causes of catheter-associated bloodstream and urinary tract infections in clinical settings (3).

Previous comparative genomic studies revealed that subspecies exist within *E. faecium* (4-7). Different names have been used by different groups to describe these clades; in this study, we use the Clade A/B nomenclature. Generally speaking, MDR *E. faecium* belong to Clade A, while Clade B consists of drug-susceptible fecal commensal *E. faecium* (8). Clade A is further split into two subclades, A1 and A2, with hospital-endemic strains generally clustering in Clade A1 and sporadic infection isolates and animal isolates generally clustering in Clade A2 (8). Specific phenotypes and genomic features are enriched in Clade A1 isolates relative to Clade A2 and B isolates (8). Specifically, Clade A1 isolates have significantly higher mutation rates, larger overall genome sizes including a larger core genome, and possess more mobile elements. On the other hand, Clade A2 possesses a larger pan-genome than Clade A1 and B, possibly reflective of the broader host origins of these strains. Given that Clade A and Clade B strains would be expected to co-mingle in certain environments (for example, in hospital and municipal sewage), the phylogenetic separation among the *E. faecium* clades suggests that they are not sharing genetic information freely because of endogenous barriers to genetic exchange.

Horizontal gene transfer (HGT) is the exchange of genetic material between cells rather than the vertical inheritance of genetic material from a parental cell. Bacteria can encode genome defense mechanisms that can act in opposition to HGT. Two examples of these mechanisms are clustered regularly interspaced short palindromic repeats (CRISPR) and associated proteins (CRISPR-Cas) systems and restriction-modification (R-M) systems. CRISPR-Cas is a dynamic immune system that utilizes sequence complementarity between self (CRISPR RNAs) and foreign nucleic acid to carry out its restrictive function, whereas R-M discriminates self from foreign DNA by DNA methylation patterns. If the *E. faecium* clades encode different defense mechanisms, they may not exchange genetic information freely, thereby facilitating and maintaining phylogenetic separation. However, little is known about CRISPR-Cas and R-M in *E. faecium*. Genomic analysis suggests that these systems could contribute to the observed clade structure of *E. faecium*. For example, CRISPR-Cas systems have been identified exclusively in Clade B *E. faecium* and in sporadic Clade A-Clade B recombinant strains (8). For R-M, a predicted methyl-directed restriction endonuclease (REase) is enriched in Clade A2 and B *E. faecium* genomes relative to Clade A1 genomes (8).

Here, we focused on R-M systems and their roles in regulating gene exchange in *E. faecium* because little is known about R-M defense in this species. Moreover, there is precedent in the literature for R-M systems contributing to bacterial clade structure, as has been observed in *Burkholderia* (9) and *Neisseria* (10). Our overarching hypothesis is that the *E. faecium* clades encode different R-M systems, thereby inhibiting genetic exchange between them. In general, R-M systems are composed of cognate methyltransferase (MTase) and REase activities and are classified into different types based on the specific number and types of enzymes in the system, as well as characteristics such as methylation type and pattern, cofactor requirement, and restriction activity (11). A MTase recognizes specific sequences in the bacterial genome and transfers a methyl group to either an adenine or a cytosine, resulting in 6-methyladenine (m6A), 4-methylcytosine (m4C), or 5-methylcytosine (m5C). A REase may recognize the same sequence as a MTase and cleave that region if the sequence is unmethylated (or in some cases, if methylated). With the activities of MTases and REases, bacteria can use R-M to impede entry of non-self DNA.

In this study, we used single-molecule real-time (SMRT) sequencing and whole genome bisulfite sequencing to characterize the methylomes of representative *E. faecium* strains from Clade A1 (*E. faecium* 1,231,502; or Efm502) and Clade B (*E. faecium* 1,141,733; or Efm733). Two unique m6A methylation patterns were identified, one in each strain. These patterns were asymmetric and bipartite, which is characteristic of Type I R-M methylation motifs (12). Bioinformatic analyses were performed to identify candidate genes responsible for the methylation. A unique Type I R-M system is encoded by each strain. We have named these systems Efa502I (for Efm502) and Efa733I (for Efm733). Expression of these candidate systems in *E. faecalis* heterologous hosts followed by SMRT sequencing confirmed that they are responsible for the methylation patterns observed in Efm502 and Efm733. A functional analysis was performed in order to assess the abilities of these systems to reduce *E. faecium* HGT by transformation. In a comparative analysis among 73 *E. faecium* genomes, we found that Efa502I is significantly enriched among Clade A1 isolates, while the Type I R-M system of Efm733 appears to be strain-specific. Overall, this study is a first step towards understanding the role of R-M in regulating HGT in *E. faecium* and the potential for R-M as one mechanism for the clade structure of *E. faecium*.

## METHODS

### Bacterial strains and growth conditions

The strains used in this study are shown in Table 1. All enterococci were grown in Brain Heart Infusion (BHI) broth or agar at 37°C, unless otherwise stated. *Escherichia coli* strains were grown in Luria Broth (LB) at 37°C and with shaking at 225 rpm unless otherwise stated. Antibiotic concentrations for enterococcal strains were as follows: rifampin, 50 μg/mL; fusidic acid, 25 μg/mL; spectinomycin, 500 μg/mL; streptomycin, 500 μg/mL; chloramphenicol, 15 μg/mL. Antibiotic concentrations for *E. coli* strains were as follows: chloramphenicol, 15 μg/mL; ampicillin, 100 μg/mL. All REases were purchased from New England Biolabs (NEB) and used per the manufacturer’s instructions. PCR was performed using Taq polymerase (NEB) or Phusion (Fisher). Sanger sequencing to validate all genetic constructs was performed at the Massachusetts General Hospital DNA Core facility (Boston, MA).

**Table 1.** Strains used in this study.

| Strain | Description | Reference |
| --- | --- | --- |
| *Enterococcus faecium* |  |  |
| 1,231,502 | Clade A1 isolate; also referred to as Efm502 | (14) |
| 1,230,933 | Clade A1 isolate | (14) |
| 1,231,410 | Clade A1 isolate | (14) |
| 1,231,408 | Hybrid Clade A1/B isolate | (14) |
| 1,231,501 | Clade A2 isolate | (14) |
| 1,141,733 | Clade B isolate; also referred to as Efm733 | (14) |
| Com12 | Clade B isolate | (14) |
| Com15 | Clade B isolate | (14) |
| Efm733ΔRM | Efm733 with deletion of Efa733I (EFSG_05027-29) | This study |
| Efm502ΔRM | Efm502 with deletion of Efa502I (EFQG_01130-32) | This study |
| *Enterococcus faecalis* |  |  |
| OG1RF | Rifampicin- and fusidic acid-resistant derivative of *E. faecalis* OG1 | (46) |
| OG1SSp | Streptomycin- and spectinomycin-resistant derivative of *E. faecalis* OG1 | (46) |
| OG1RF::*efa502I* | OG1RF with Efa502I inserted at GISE site | This study |
| OG1SSp::*efa733I* | OG1SSp with Efa733I inserted at GISE site | This study |
| *Escherichia coli* |  |  |
| EC1000 | Plasmid propagation host | (20) |
| DH5α | Plasmid propagation host |  |
| STBL4 | Plasmid propagation host | Fisher |
| STBL4(pGEM-T) | STBL4 with pGEM-T Easy | This study |
| STBL4(pRB01) | STBL4 with pGEM-T Easy vector containing EFSG_00659 | This study |
| Plasmids |  |  |
| pGEM-T Easy | Commercial plasmid for gene propagation | Promega |
| pLT06 | Temperature-sensitive plasmid | (19) |
| pAT28 | Shuttle vector for *E. faecalis* | (22) |
| pWH03 | Used for gene insertion in the enterococci | (16) |
| pRB01 | pGEM-T Easy Vector with EFSG_00659 | This study |
| pHA102 | pWH03 containing NotI-digested fragments of Efa733I M and S loci (EFSG_05028-9) | This study |
| pHA103 | pWH03 containing NotI-digested fragments of Efa502I M and S loci (EFQG_01131-2) | This study |
| pWH16 | pLT06 containing fragments from upstream and downstream of Efa502I | This study |
| pWH17 | pLT06 containing fragments from upstream and downstream of Efa733I | This study |

### Isolation of genomic DNA

Enterococcal strains were cultured overnight in BHI broth prior to genomic DNA (gDNA) extraction. The extraction was performed using a Qiagen Blood and Tissue DNeasy Kit using a previously published protocol (13). To isolate *E. coli* gDNA, bacteria were grown overnight in LB broth prior to extraction using either the Blood and Tissue DNeasy kit (Qiagen) or the UltraClean Microbial DNA Isolation Kit (Qiagen) per the manufacturer’s instructions. Whole genome amplification (WGA) control DNA was generated by amplification of native gDNA using the REPLI-g kit (Qiagen) per the manufacturer’s instructions.

### SMRT sequencing and methylome detection in *E. faecium*

SMRT sequencing of *E. faecium* gDNA and their WGA controls was performed by the Johns Hopkins Medical Institute Deep Sequencing and Microarray Core. After sequencing, the reads were aligned to existing references (for Efm502, NZ_GG688486-NZ_GG688546 and for Efm733, NZ_GG688461-NZ_GG688485) and analyzed using the RS modification and motif detection protocol in SMRT portal v1.3.3. WGA controls were used as methylation baselines.

### Bioinformatic analysis of R-M systems in eight *E. faecium* genomes

The entire protein complement for eight previously sequenced *E. faecium* isolates (14) was analyzed. To identify potential MTases, the REBASE Gold Standard list (15) was used as a reference. This list is comprised of biochemically verified MTases and REases. Each protein sequence from *E. faecium* genomes was analyzed using BlastP against the REBASE Gold Standard list. The protein sequences with significant (e-value <1e^-3^) homology to REBASE Gold Standard proteins were further filtered by protein size. If an *E. faecium* query protein length was less than half of its subject’s length, the match was removed from the prediction list. Due to the sequence diversity of REases which complicates their bioinformatic identification (15), guilt-by-association was used to identify full R-M systems as we have previously described (16). The proteins encoded near candidate DNA MTases were analyzed using BLAST and Pfam for conserved domains consistent with REase activities and/or sequence identity to confirmed REases. The amino acid sequence of each R-M candidate was then pairwise compared among all the eight strains to identify putative orthologs. If two protein sequences shared an amino acid identity ≥90% with query coverage ≥90%, they were considered to be orthologous.

### Expression of R-M systems in *E. faecalis* heterologous hosts

Genes encoding the specificity and methylation subunits of Efa733I (EFSG_05028-EFSG_05027) were PCR-amplified in their entirety, including the upstream region to retain the native promoter, using primers 733_T1A_SM_F and 733_T1A_SM_R (see Table 2 for primer sequences). The PCR product was digested with *Not*I and ligated into *Not*I-digested pWH03 (16) using T4 DNA Ligase (NEB), generating pHA102. pWH03 is a pLT06 derivative for expression of genes from a previously validated neutral genomic insertion site (EF2238-EF2239) for expression (GISE) (16, 17). pHA102 constructs were then introduced into *E. coli* DH5α via heat shock for propagation and sequence confirmation. pHA102 was electroporated into *E. faecalis* OG1SSp using a previously described method (18). An *E. faecalis* OG1SSp derivative with a chromosomal integration of Efa733I, referred to as OG1SSp::*efa733I*, was generated by temperature shifts and *p-*chlorophenylalanine counterselection, as previously described (19).

**Table 2.**
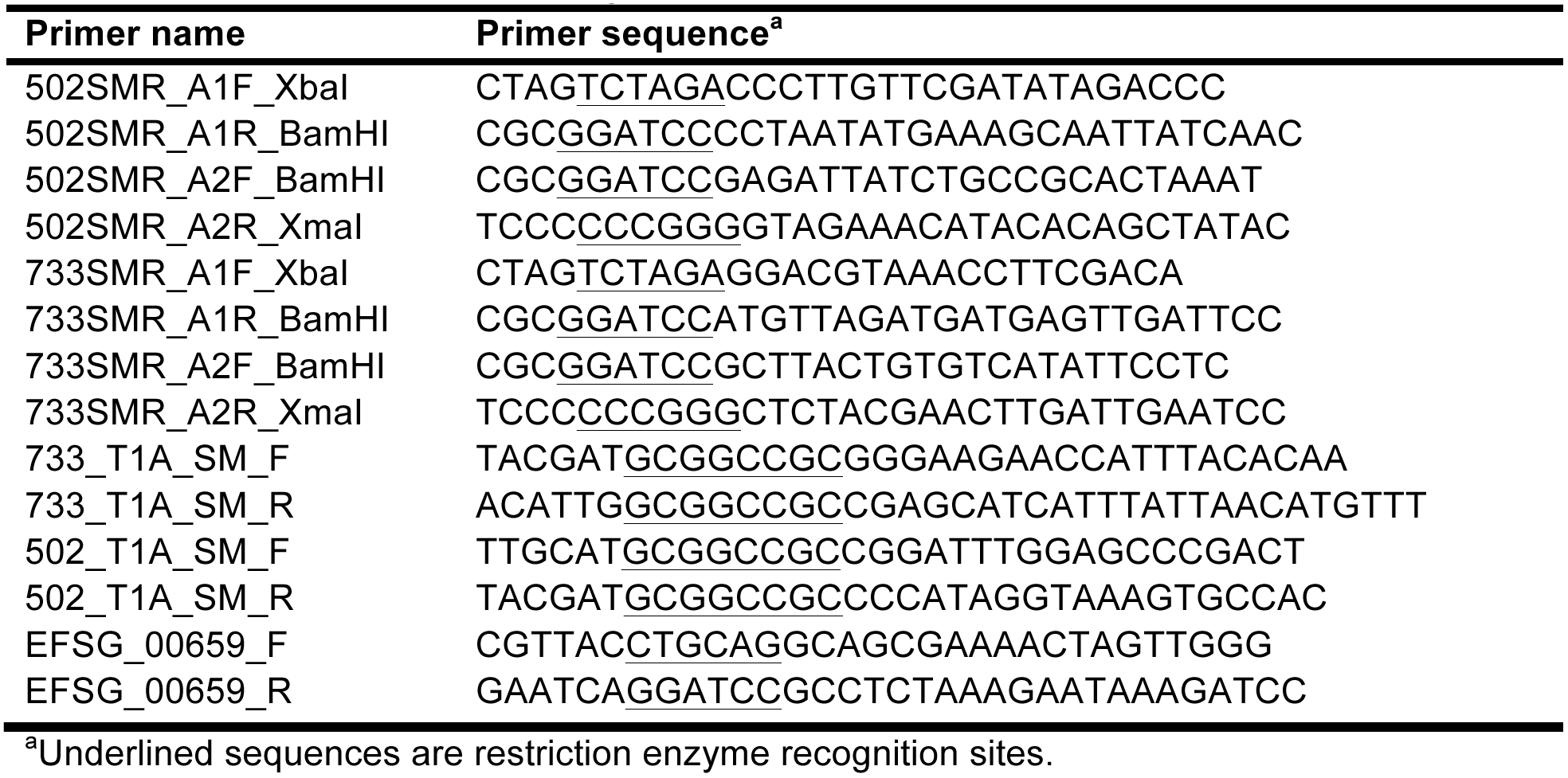
Primers used in this study.

Genes encoding the specificity and methylation subunits of Efa502I (EFQG_01131-EFQG_01132) were PCR-amplified using primers 502_T1A_SM_F and 502_T1A_SM_R. The PCR product was then TA-cloned into the pGEM-T Easy Vector (Promega) and introduced into DH5α via heat shock to generate pGEM-SMA1. pGEM-SMA1 was then digested with *Not*I, and the digestion reaction was used as insert for ligation into *Not*I-digested pWH03. The ligation reaction was then introduced into DH5α, and colonies were screened for chloramphenicol resistance and ampicillin susceptibility to ensure the pGEM backbone was not ligated into pWH03. Once the construct, referred to as pHA103, was confirmed via Sanger sequencing, it was introduced into electrocompetent *E. faecalis* OG1RF. An *E. faecalis* OG1RF derivative with a chromosomal integration of Efa502I, referred to as OG1RF::*efa502I*, was generated by temperature shifts and *p-*chlorophenylalanine counterselection. All plasmids and strains for heterologous expression were validated by PCR and Sanger sequencing.

SMRT sequencing in *E. faecalis* OG1 derivatives expressing Efa502I or Efa733I was performed by the University of Michigan sequencing core facility. Reads were mapped to the *E. faecalis* OG1RF reference sequence (GenBank accession number NC_017316), and the methylation motifs were detected using the RS modification and motif detection protocol in SMRT portal v.2.3.2. *In silico* controls were used as modification baselines.

### Generation of *E. faecium* R-M deletion mutants

Regions up- and downstream of Efa502I and Efa733I were PCR-amplified using primers listed in Table 2, ligated into pLT06, and transformed into *E. coli* EC1000 (20), generating pWH16 and pWH17 (Table 1). Insert sequences were confirmed using Sanger sequencing. *E. faecium* strains were made electrocompetent using previously a published protocol (21). 2 μg of sequence-confirmed plasmids were electroporated into electrocompetent Efm733 and Efm502. The generation of deletion mutants was accomplished using temperature shifts and *p-*chlorophenylalanine counterselection, as previously described (19). The successful deletion mutants were sequence-confirmed by PCR and Sanger sequencing.

### Transformation efficiency test

Efm733, Efm502, and their respective R-M deletion mutants were made electrocompetent using a modified version of the previously published protocol (21). Briefly, overnight cultures were diluted 10-fold in BHI and cultured to OD_600nm_ ~ 0.6. The bacteria were then pelleted and treated with filter-sterilized lysozyme buffer (10 mM Tris-HCl pH 8.0, 10 mM EDTA pH 8.0, 50 mM NaCl) supplemented with 83 μL of 2.5 KU/mL mutanolysin stock for 30 min at 37°C. The cells were then pelleted and washed three times with ice-cold filter-sterilized electroporation buffer (0.5 M sucrose and 10% glycerol). Finally, the cells were pelleted and resuspended in electroporation buffer and aliquoted for storage at −80°C and future use. 1 μg pAT28 (22) was electroporated into the electrocompetent *E. faecium* cells. The counts of total viable cells and spectinomycin-resistant cells were determined by serial dilution and plating. The transformation efficiency was expressed as percent of transformed (spectinomycin-resistant) cells per total viable cells. Three independent experiments were performed and the statistical significance was assessed using the unpaired one-tailed Student’s t-test.

### Distribution analysis of putative R-M systems and orphan MTases

The amino acid sequences for select R-M system and orphan MTase candidates were queried against a collection of 73 *E. faecium* isolates previously analyzed by Lebreton *et al* (8) using BLASTP. Any proteins which shared >90% query coverage and amino acid identity were considered orthologs. The Fisher’s exact test was used to determine if an orphan MTase or R-M system was significantly over- or under-represented in a particular clade.

### REase protection assays

To identify m5C methylation, gDNA was treated with the methylation-sensitive REases McrBC, FspEI, and MspJI (NEB). 500 ng gDNA was incubated with each REase at 37°C for 3 h (McrBC) or 6 h (FspEI and MspJI) followed by analysis by electrophoresis on a 1% agarose gel with ethidium bromide.

### Bisulfite sequencing

Whole-genome bisulfite sequencing libraries were constructed using the Illumina TruSeq LT PCR FREE kit and the Qiagen EpiTect Bisulfite kit. Native DNA was isolated as described above. Whole-genome-amplified (WGA) control DNA was generated by amplification of native gDNA using the Qiagen REPLI-g^®^ kit, per the manufacturer’s instructions. For bisulfite sequencing, briefly, 2 μg each of native and WGA control DNA were fragmented using NEB fragmentase. DNA fragments ranging from 200 bp to 700 bp were gel extracted and end-repaired. After A-tailing of DNA fragments, Illumina TruSeq adapters were ligated. Then, the bisulfite conversion was performed using the Qiagen EpiTect Bisulfite kit, per the manufacturer’s instruction. An 8-cycle PCR enrichment with Illumina primer mix was performed, followed by size selection and gel purification. The libraries were sequenced using Illumina MiSeq with 2×75 bp paired-end chemistry.

### Whole genome bisulfite sequencing analysis

The sequencing reads were analyzed using Bismark (23) with additional quality control and filtering as described previously (16). Briefly, the Illumina reads were mapped to the *in silico* bisulfite-converted references (23). Then, we quantified the conversion rate of each mapped read by calculating the percentage of converted C (which will result in T) to the total number of C in the reference within the mapped region. The mapped reads with ≤80% conversion rate were filtered out from analysis (16). Next, the coverage depth and methylation ratio were calculated for each C site. The methylation ratio was calculated by dividing the total number of C by the coverage depth at each C site. A fully methylated C, thus protected from bisulfite conversion, will have a methylation ratio near 1. An unmethylated C will have a methylation ratio near 0. To identify consensus methylation motifs, C sites with ≥0.35 methylation ratio and ≥10 coverage depth, along with the sequences of 5 bp upstream and 5 bp downstream, were extracted. The extracted sequences were subjected for MEME motif search (24).

### Confirmation of m5C MTase activity

Primers EFSG_00659_F and EFSG_00659_R (Table 2) were used to amplify the entire Efm733 EFSG_00659 coding region and its upstream predicted promoter. The PCR product was then cloned into the pGEM-T Easy Vector (Promega) per the manufacturer’s instructions and transformed into *E. coli* STBL4 (Fisher) to generate pRB01. REase digestion assays with methylation-sensitive enzymes were performed on purified *E. coli* and *E. faecium* gDNA as described above.

### Accession numbers

DNA sequence data generated in this study have been deposited in the Sequence Read Archive under accession numbers PRJNA397049 (for SMRT sequencing data) and PRJNA488088 (for Illumina bisulfite sequencing data).

## RESULTS

### Identification of Clade A1-specific putative Type I R-M system in *E. faecium*

We previously reported that a Type II R-M system significantly reduces HGT via conjugation (18) and transformation (16) in *E. faecalis*. Here, we hypothesize that the *E. faecium* clades encode distinct R-M systems that reduce the exchange of genetic information between them. We utilized an approach we previously developed for *E. faecalis* R-M analysis (18) to predict potential R-M systems in eight previously sequenced *E. faecium* genomes. The 8 genomes included 3 genomes from Clade A1, 3 genomes from Clade B, one genome from Clade A2, and one recombinant Clade A1/B hybrid (5, 8). Because REases are difficult to identify with bioinformatics, and MTase prediction is comparatively straightforward, as has been previously reported by NEB (15), we first identified predicted DNA MTases in *E. faecium* genomes, and then analyzed surrounding genes for predicted R-M-related activities. The complete list of candidates for the eight strains is shown in Table 3.

**Table 3.** Distribution of predicted DNA MTases and R-M Systems*.

| Strain name |  | 1,230,933 | 1,231,410 | 1,231,502 | 1,231,501 | 1,231,408 | 1,141,733 | Com12 | Com15 |
| --- | --- | --- | --- | --- | --- | --- | --- | --- | --- |
| Clade |  | A1 | A1 | A1 | A2 | Hybrid A1/B | B | B | B |
| Type I Restriction Modification System | Specificity | EFPG_01270 | EFTG_00783 | EFQG_01131 |  | EFUG_01527 | EFSG_05028 |  | EFWG_02518 |
|  | Modification | EFPG_01269 | EFTG_00782 | EFQG_01132 |  | EFUG_01526 | EFSG_05029 |  | EFWG_02519 |
|  | Restriction | EFPG_01271 | EFTG_00784 | EFQG_01130 |  | EFUG_01528 | EFSG_05027 |  | EFWG_02517 |
| Type I Restriction Modification System | Specificity | EFPG_05435 and EFPG_05434 | EFTG_02031 |  |  |  |  |  |  |
|  | Modification | EFPG_05433 | EFTG_02032 |  |  |  |  |  |  |
|  | Restriction | EFPG_05436 | EFTG_02030 |  |  |  |  |  |  |
| Type I Restriction Modification System | Specificity |  |  |  | EFRG_00106 |  |  |  |  |
|  | Modification |  |  |  | EFRG_00107 |  |  |  |  |
|  | Restriction |  |  |  | EFRG_00105 |  |  |  |  |
| Type II Methyltransferase |  | EFPG_01821 | EFTG_02364; EFTG_02653 | EFQG_01609, EFQG_02270 |  |  | EFSG_00659 |  |  |
| Unspecified Methyltransferases |  |  | EFTG_02333 |  | EFRG_02239 | EFUG_01512 |  | EFVG_00502 |  |

\*Loci that are in black are strain specific. Loci that are the same color are conserved in their protein sequences based on a >90% sequence identity threshold. An empty cell indicates that the system was not detected.

Interestingly, we predicted at least one putative Type I R-M system for seven of the eight *E. faecium* strains (Table 3). Type I R-M systems are multisubunit complexes comprised of a specificity subunit (S), a methylation subunit (M), and a restriction subunit (R) (11, 25-27). The S subunit is responsible for the specific DNA recognition motif and associates with the DNA to bring the M and R subunits into contact. The system has two conformations: M_2_S_1_, which is capable of methylating DNA based on the recognition sequence, and R_2_M_2_S_1_, which is capable of restricting DNA (27, 28). One predicted *E. faecium* Type I R-M system is comprised of highly conserved (>90% amino acid sequence identity) M and R subunits in six of eight genomes across both Clade A1 and Clade B (Table 3 and Dataset S1). The specificity subunit from this system, however, is highly conserved in Clade A1 genomes but not in Clade B (Table 3; Fig. 1 and Fig S1). S subunits possess two target recognition domains (TRDs) that determine the nucleotide sequence the subunit binds to (29, 30). The variation in amino acid sequence between the S subunits occurs within these TRDs (Fig S1), suggesting that these S subunits recognize different DNA sequences. Notably, the S subunits from Clade A1 strains are identical to each other, indicating that they utilize the same recognition sequence.

**Figure 1.**
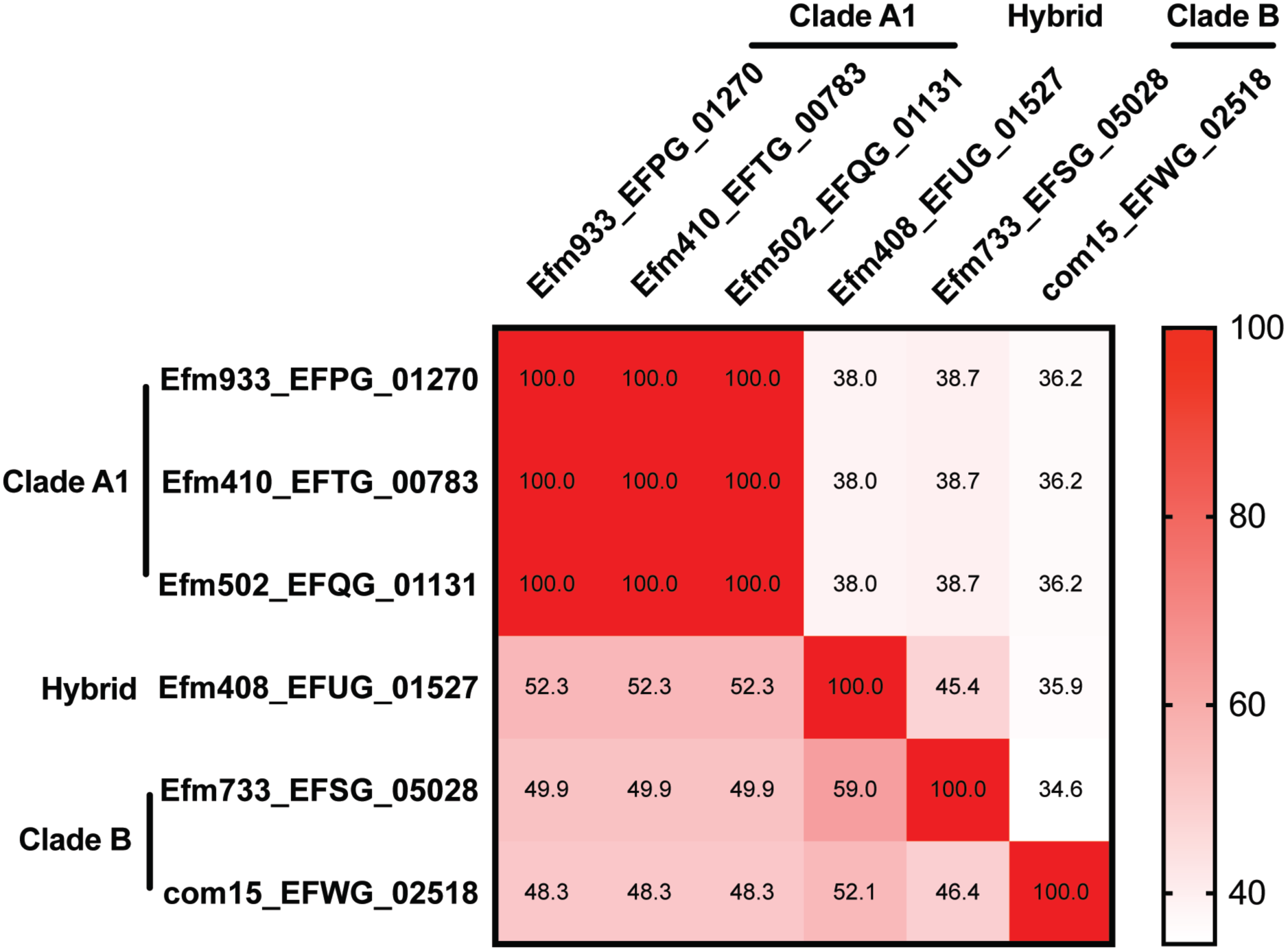
Conservation and variability of S subunits. The protein sequences of predicted S subunits from 6 (out of 8) representative *E. faecium* genomes were pairwise aligned using MacVector. The percent identity of each pair is shown. White to red: low to high percent identities. Each number represents percent identity of one protein sequence (row name) to another (column name).

To examine the distribution of this putative Clade A1-specific system in a larger collection of *E. faecium* strains, we analyzed 73 *E. faecium* genomes of mostly draft status that were reported previously (8). This list includes 15 clade B isolates, 21 clade A1 isolates, 35 clade A2 isolates, and 2 hybrid isolates (Table S1). We selected Efm502 (Clade A1) as our representative Clade A1 strain for this analysis and used Type I R-M sequences from this genome as references for analysis against the broader collection of *E. faecium* strains. The M and R subunits of the putative Clade A1-specific Type I system were detected in 51 and 52, respectively, of 73 *E. faecium* genomes, including both Clade A and Clade B strains (Fig S2a). However, the distribution of the S subunits varied (Fig S2a-b). The S subunit present in Efm502, EFQG_01131, was significantly enriched within Clade A1 isolates (14/21; p-value <0.0001 using Fisher exact test; Fig S2a) and absent from all other clades with the exception of strain EnGen002, which is classified as a Clade A1/B hybrid strain. Interestingly, the S subunits present in most other *E. faecium* strains are strain-specific by the strict thresholds applied here. Given that the Efm502 S subunit is enriched in Clade A1, we hypothesize that many Clade A1 strains exchange genetic information freely with each other while exchange with other *E. faecium* strains is restricted.

### SMRT sequencing for *E. faecium* methylome analysis

We analyzed the Efm502 and Efm733 genomes by SMRT sequencing. SMRT sequencing measures the kinetics of DNA polymerase as it synthesizes DNA in order to identify bases that have been modified (31-33). It has been extensively utilized for bacterial methylome analysis (34-42). With SMRT sequencing, 6-methyladenine (m6A) and 4-methylcytosine (m4C) can be easily detected with modest sequence coverage (~25x per strand), while 5-methylcytosine (m5C) detections requires high coverage (~250x per strand) (41, 43). Using SMRT sequencing, we identified two unique m6A methylation motifs in Efm502 and Efm733. Efm502 possessed m6A methylation at the underlined position of the motif 5’-R<u>A</u>YCNNNNNNTTRG-3’ (and its complementary strand 5’-CYA<u>A</u>NNNNNNGRTY-3’) and Efm733 possessed m6A methylation at the underlined position of the motif 5’-<u>A</u>GAWNNNNATTA-3’ (and its complementary strand 5’-TA<u>A</u>TNNNNWTCT-3’) (Table 4; Dataset S2). These sequences are asymmetric and bipartite, which is characteristic of Type I R-M methylation (12). Due to the coverage of our SMRT sequencing, m5C modification could not be accurately detected. The two unique m6A methylation patterns indicate that DNA from one strain would be recognized as foreign should it cross the strain barrier.

**Table 4.**
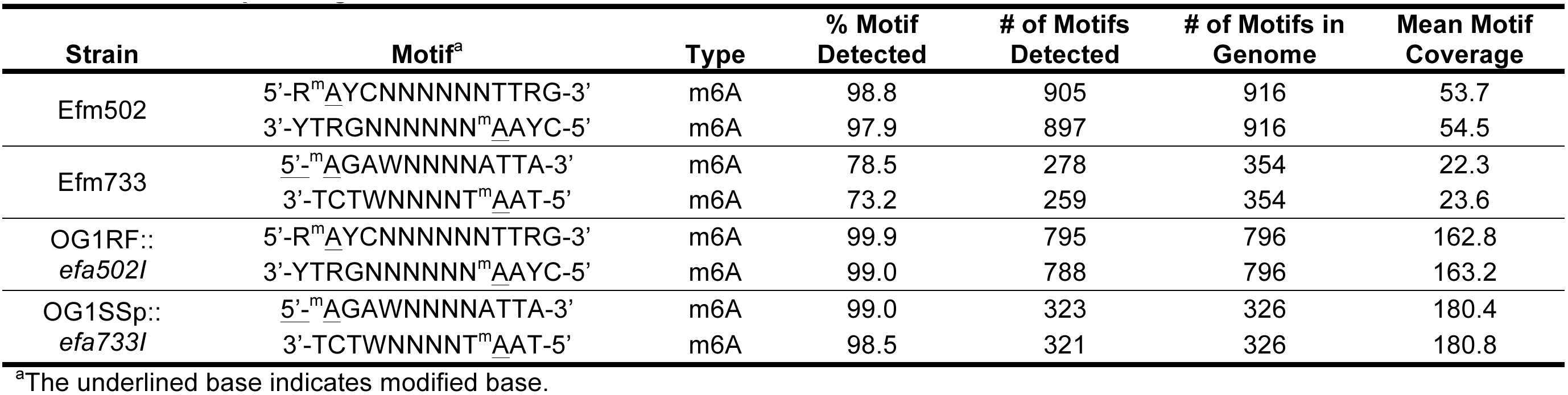
SMRT Sequencing results.

### Expression in heterologous hosts links methylation activity to genes in Efm502 and Efm733

According to our predictions (Table 3), there is only one complete Type I R-M system encoded by each of Efm502 and Efm733. To determine if these systems are responsible for the methylation patterns identified by SMRT sequencing, we expressed the respective S and M subunits (EFQG_01131-01132 for Efm502 and EFSG_05028-05027 for Efm733) in the heterologous host *E. faecalis* OG1RF or its spectinomycin/streptomycin-resistant relative OG1SSp. Previous work in our lab had characterized the methylome of OG1RF using SMRT and bisulfite sequencing (16). This allowed us to attribute any new methylation patterns observed during SMRT sequencing to the *E. faecium* genes that were expressed in the OG1RF background. SMRT sequencing of these strains detected the same methylation patterns originally identified in Efm502 and Efm733 (Table 4; Dataset S2). These data demonstrate that EFQG_01131-01132 is responsible for the 5’-R<u>A</u>YCNNNNNNTTRG-3’ methylation in Efm502 and that EFSG_05028-05027 is responsible for the 5’-<u>A</u>GAWNNNNATTA-3’ methylation in

Efm733. Because we have confirmed the function of these genes, we have named them Efa502I and Efa733I, which is consistent with the R-M system nomenclature convention established by New England Biolabs (12).

### Efa502I and Efa733I reduce transformation efficiency in *E. faecium*

To determine whether the Type I R-M systems in Efm733 and Efm502 actively defend against exogenous DNA, we constructed null strains (Efm733∆RM and Efm502∆RM; Table 1) and evaluated their transformation efficiencies relative to their wild-type parent strains. Here, we utilized the broad host range plasmid pAT28 (Table 1) (22). pAT28 sequence has motifs recognized by the Type I R-M systems in both wild-type Efm733 (1 occurrence) and Efm502 (1 occurrence). The transformation of pAT28 into Efm733 and Efm502 served as a baseline for the experiment. If Efa733I and Efa502I are active, we expect to see higher pAT28 transformation efficiencies into Efm733∆RM and Efm502∆RM, respectively. Indeed, we observed significantly higher transformation efficiencies into Type I R-M system null strains (Fig 2; p-value<0.05 using one-tailed Student’s t-test). These data demonstrate that the Type I R-M systems in Efm733 and Efm502 actively function as mechanisms of genome defense.

**Figure 2.**
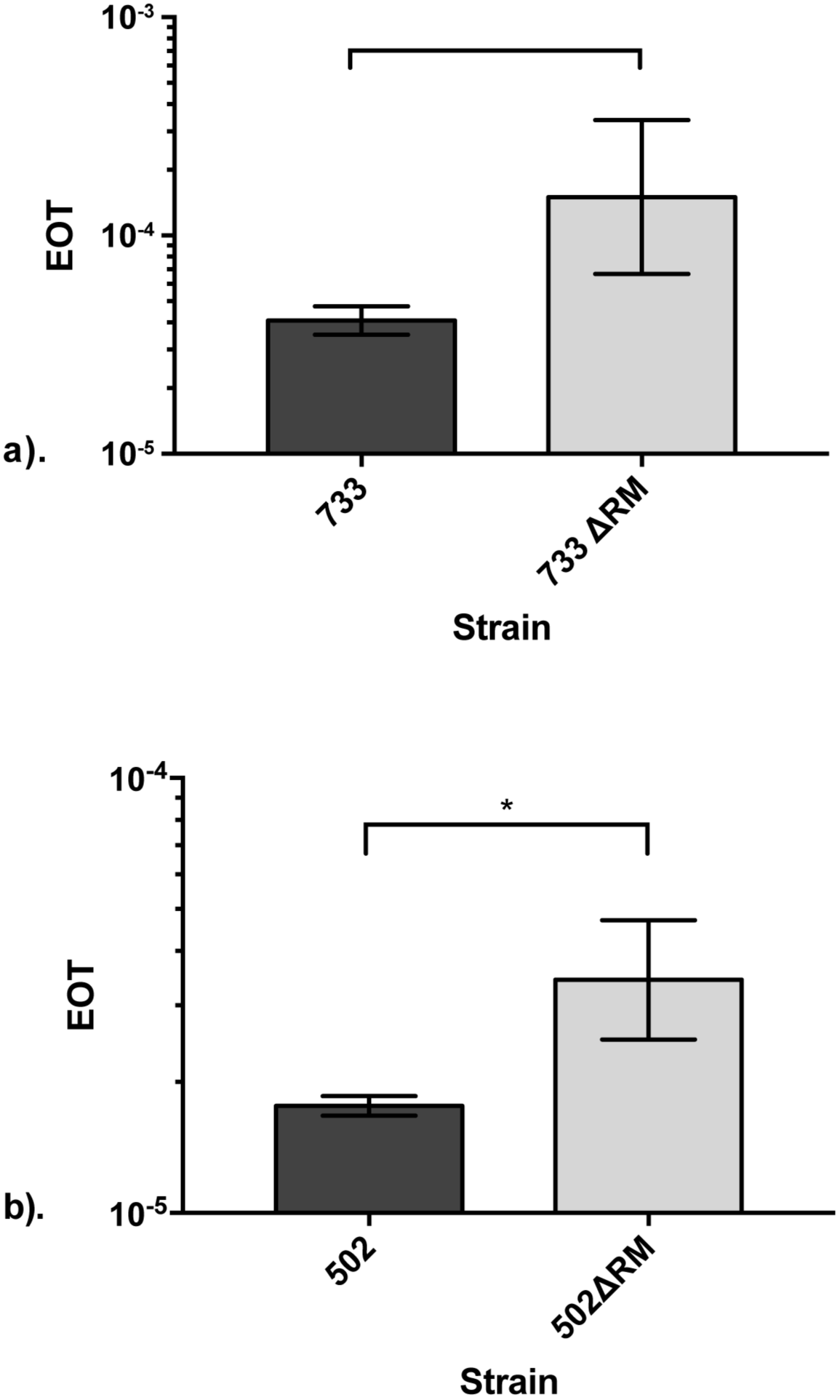
Type I R-M systems reduce transformability of Efm733 and Efm502. Three independent transformation experiments were performed. There is a statistical difference between the transformation efficiency of pAT28 into wild type and R-M null strains of Efm733 and Efm502. EOT: Efficiency of transfer. *: p < 0.05.

### m5C methylation occurs in Efm733

As described previously, our SMRT sequencing had insufficient coverage depth for m5C methylome characterization. Hence, we used REase protection assays with commercially available methylation-sensitive REases to query the presence of m5C methylation in our eight *E. faecium* strains. Table 5 summarizes the recognition sequences and modifications of the enzymes used in this study. Only Efm733 showed evidence of cytosine modification, as it was digested by MspJI (Table 5; Fig S3).

**Table 5.**
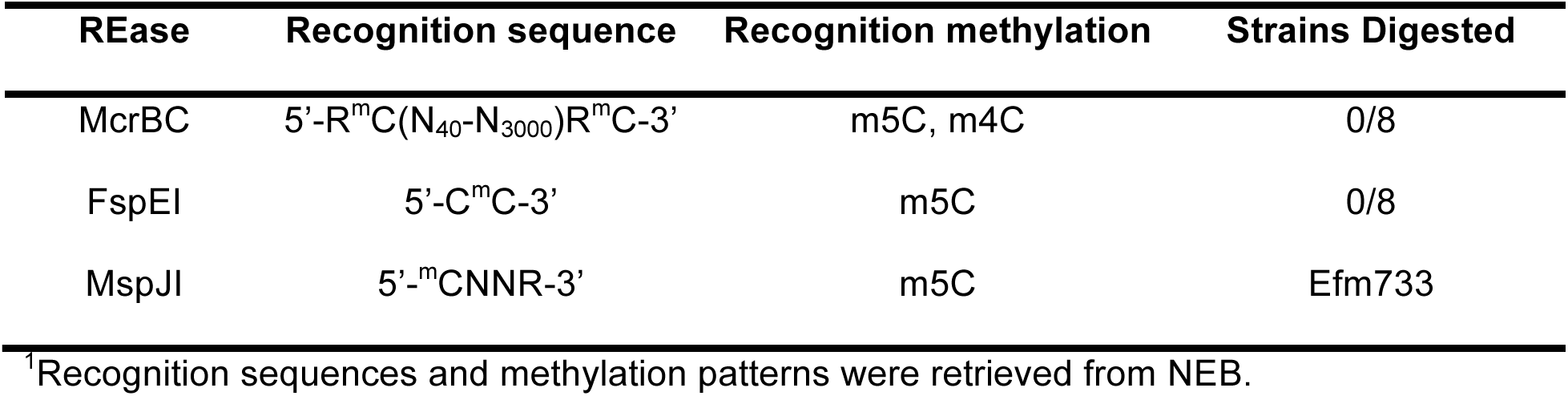
Methylation-sensitive REase digestion reaction results^1^.

To determine the exact cytosine methylation motif present in Efm733, gDNA was subjected to whole-genome bisulfite sequencing. Whole genome amplified (WGA) DNA was used as negative control since WGA removes all modifications. During bisulfite treatment, cytosine bases are converted to thymine unless they are protected by either m4C or m5C methylation. Additionally, our lab has previously published a method of distinguishing between m4C and m5C methylation using thymine conversion ratios of sequencing reads after bisulfite treatment (44). m5C methylation is sufficient to protect the cytosine residue completely from bisulfite conversion, so that most sequencing reads at that position contain the original cytosine base. However, m4C methylation provides only partial protection from bisulfite conversion, so a thymine conversion rate of 0.5 at a particular position within the sequencing reads suggests the presence of m4C methylation. Bisulfite conversion and subsequent sequencing revealed that Efm733 possesses m5C modification at the motif 5’-R^m^<u>C</u>CGGY-3’ (Table 6, Fig 3, and Fig S4; the methylation occurs at the underlined position), which overlaps the MspJI recognition site (5’-CNNR-3’) and hence supports the evidence of methylation obtained from the MspJI digestion assays. Based on the cytosine conversion ratio of close to 1.0, m5C modification is supported, which is consistent with why it was not detected by our SMRT sequencing.

**Figure 3.**
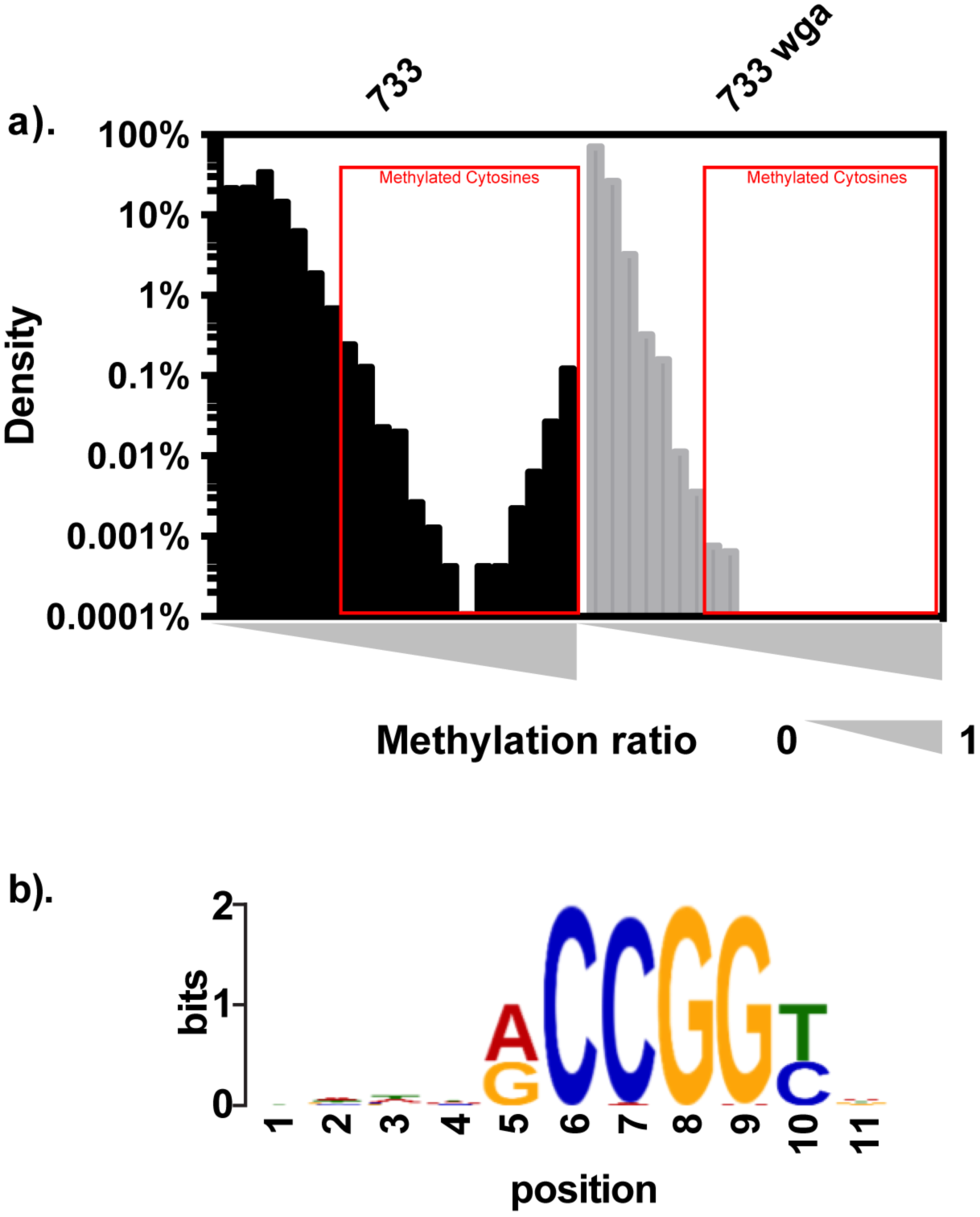
Bisulfite sequencing results for Efm733. Efm733 gDNA and its whole genome amplified (WGA) control DNA were bisulfite-converted and deep sequenced. The methylation ratio for each cytosine site was calculated. A). The methylation ratio was plotted against the density of cytosine sites with that methylation ratio. The presence of cytosine sites with methylation ratio near 100% in Efm733 native gDNA but not WGA control indicates the presence of m5C methylation. All cytosine sites with ≥0.35 methylation ratio as indicated with red boxes from native gDNA samples were extracted, together with 5 bp upstream and 5 bp downstream sequences. The sequences are subjected to consensus motif analysis using MEME. The consensus motif identified by this analysis is shown in (b) and the center position (position 6) indicates where the modification was detected.

**Table 6.**
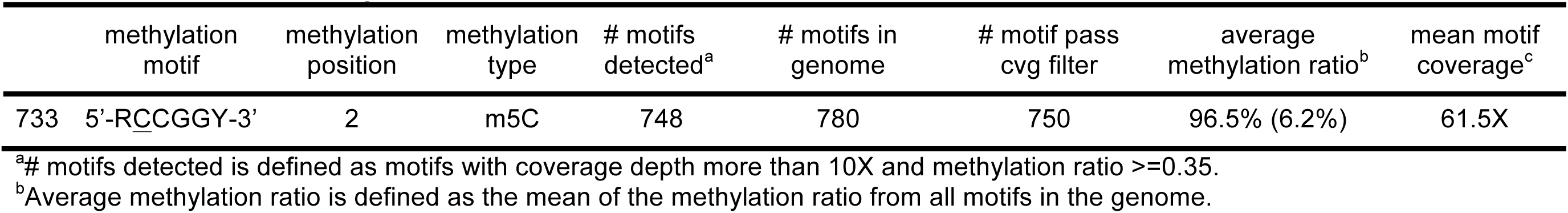
Bisulfite sequencing results.

### EFSG_00659 is responsible for m5C methylation in Efm733

Based on the bioinformatics analyses, we hypothesized that EFSG_00659 was responsible for the m5C methylation found in Efm733 (Table 3). EFSG_00659 possessed no homologs in the other seven strains analyzed, making it a good candidate for the unique methylation found in Efm733. We queried the EFSG_00659 protein sequence against the REBASE gold standard list and identified that it has high sequence similarity to M.AvaIX, M.VchO395I and M.VchAI (e-value ≤ 3e^-125^; recognition sites are 5’-RCCGGY-3’). Interestingly, BLASTP identified no significant hits when EFSG_00659 was queried against the larger collection of 73 *E. faecium* genomes, indicating its unique presence in Efm733.

In order to link EFSG_00659 with the m5C methylation identified during bisulfite sequencing, we expressed it in the heterologous host *E. coli* STBL4 and performed an REase protection assay. The REase AgeI recognizes the motif 5’-A<u>C</u>CGGT-3’, and its enzymatic activity is blocked if m5C methylation is present at the underlined position. This motif overlaps the m5C methylation motif in Efm733 identified by bisulfite sequencing. If the motif is methylated, DNA will be protected against digestion. We cloned EFSG_00659 into the vector pGEM-T and transformed it into *E. coli* STBL4, generating strain *E. coli* STBL4(pRB01). Genomic DNA from Efm733, STBL4(pGEM-T), and STBL4(pRB01) was treated with AgeI per the manufacturer’s instructions. EcoRI was used as a positive control for digestion. Figure 4 shows representative results of the digestions on a 1% agarose gel. As expected, Efm733 was digested by EcoRI and protected against digestion from AgeI. STBL4(pGEM-T) was digested by both EcoRI and AgeI, indicating that the original host and empty vector pGEM-T did not possess the appropriate m5C methylation. STBL4(pRB01) was protected against digestion by AgeI, demonstrating that EFSG_00659 is responsible for the 5’-R^m^CCGGY-3’ methylation found in Efm733.

**Figure 4.**
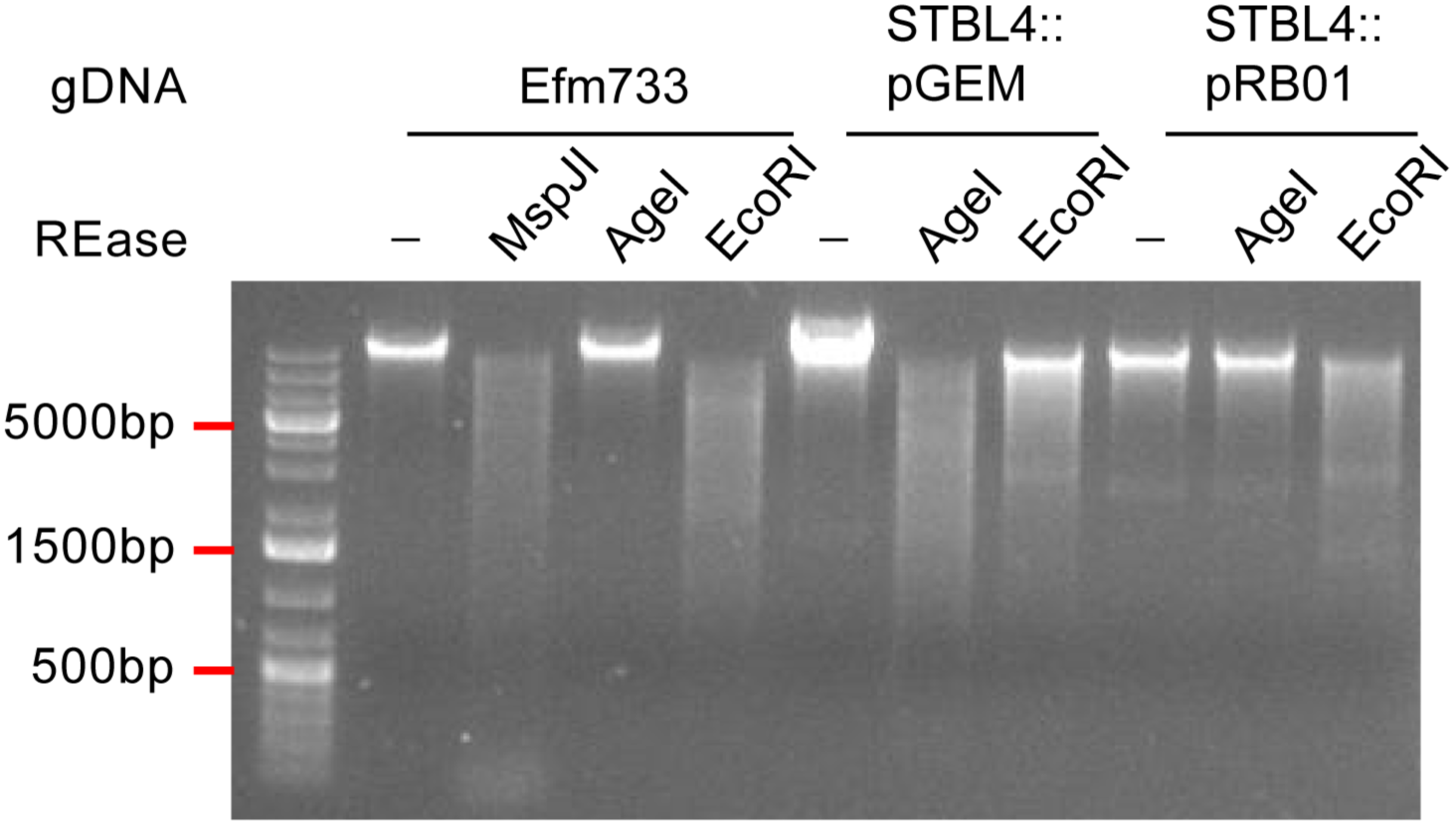
EFSG_00659 confers protection against AgeI digestion. AgeI digestion reactions were analyzed by agarose gel electrophoresis with ethidium bromide staining. Bacterial gDNA was used as substrate for REase reactions. Expression of the Efm733 gene EFSG_00659 in *E. coli* STBL4 protects *E. coli* gDNA from AgeI digestion. EcoRI is a positive control for digestion. -, no enzyme added.

## DISCUSSION

In this study, we used a combination of genomic and genetic approaches to identify a functional Type I R-M system that is enriched in Clade A1 *E. faecium* and that significantly alters transformability of a model Clade A1 strain. We propose that this R-M system impacts HGT rates among *E. faecium* mixed-clade communities, thereby helping to maintain the observed phylogenetic structure of *E. faecium* and facilitating HGT specifically among Clade A1 strains. Mixed communities of *E. faecium* clades are expected to occur in environments where healthy and ill human hosts, human and animal hosts, and/or the feces of any of these hosts co-mingle (i.e. in sewage). In future studies, we plan to assess the impact of R-M on conjugative plasmid transfer, which is a major mode of HGT in enterococci and was not assessed in our current study.

An interesting observation from our study is the sequence diversity of Type I S subunits encoded within *E. faecium* Type R-M systems having nearly identical R and M subunits (Fig S2a-b). After further investigation into those alignments, we found that those S subunits sharing 50-70% overall amino acid sequence identities possess sequence diversity within one TRD domain, where the other TRD domain and the central conserved domain are conserved. This suggests that these systems share partial recognition sequences. Previous research has reported that the diversification of Type I R-M recognition sequences is driven by TRD exchanges, permutation of the dimerization domain, and circular permutation of TRDs (45). Our observation suggests that TRD recombination and reorganization events occur for *E. faecium* Type I R-M systems outside Clade A1. Future studies will use genomics to further explore the relationship between S subunit sequence diversity and its impact on *E. faecium* methylomes and inter-strain and inter-clade HGT.

## Acknowledgments

This work was supported by Public Health Service grants K22AI099088 and R01AI116610 to K.L.P.

